# Recurrent non-canonical proteoforms in acute myeloid leukemia identified by integrative proteogenomics

**DOI:** 10.64898/2026.08.27.747547

**Authors:** Laura K. Schmalbrock, Asher Preska Steinberg, Katarzyna Kulej, Jinxin Zhang, Gabriella Casalena, Andrew McPherson, Alex Kentsis

## Abstract

Reference proteomes incompletely represent proteins translated in cancer, leaving tumor-specific proteoforms outside the search space of conventional mass spectrometry (MS). Such “dark proteome” products may arise from genomic variation, aberrant transcription or splicing, and non-canonical translation, including microproteins encoded by small open reading frames (ORFs). To define this landscape in acute myeloid leukemia (AML), we developed a cohort-informed proteogenomic strategy using paired RNA-sequencing and MS analysis of 123 human patient AML specimens and 13 healthy CD34+ controls. ProteomeGenerator2 was used for de novo transcriptome assembly and ORF prediction, and candidate cancer-specific unannotated sequences were prioritized by unique high-quality mass spectral support, absence from CD34+ controls, recurrence across individual AML patients, and lack of close homology to annotated proteins. We identified 5,849 Swiss-Prot-unannotated proteoforms, including 1,987 without homology to annotated human proteins. Thirty-nine candidates, most encoding microproteins, were recurrently detected in more than 10% of patients, and 14 were independently validated by deep, fractionated, multi-protease data-independent acquisition (DIA) proteomics of human AML cell lines. Structural modeling predicted several functional classes, including intrinsically disordered, alpha-helical microproteins, and membrane- or secretory-pathway-associated proteoforms. These findings define a recurrent AML dark proteome and establish a framework for the discovery of tumor-specific non-canonical proteins for mechanistic and therapeutic studies.

## Introduction

Mass spectrometry identifies proteins by matching peptide spectra to protein sequence databases. As a result, protein discovery is limited to annotated proteins. Curated reference proteomes such as UniProtKB/Swiss-Prot^1^ provide high-confidence maps of canonical proteins and known isoforms, but they incompletely represent the full set of proteins that may be translated in cancer. Somatic mutations, aberrant transcription, alternative splicing, and non-canonical translation can generate reference-unannotated proteoforms, including microproteins translated from small open reading frames (smORFs)^2, 3, 4, 5, 6^. Recent advances in long-read RNA sequencing^7^, ribosome profiling^8, 9, 10^, high-resolution mass spectrometry^11, 12^, and integrative proteogenomic analysis^13, 14^ now make it possible to elucidate these cryptic translated products more systematically. A central challenge is to distinguish bona fide translated proteoforms from transcriptomic predictions, database artifacts, and peptide matches to related annotated canonical proteins. Ongoing work aims across multiple fields to systematically discover such uncharacterized proteins and define their functions in health and disease.^5, 10, 15, 16, 17^

Acute myeloid leukemia (AML) remains fatal for many patients despite advances in molecular diagnosis and targeted therapy, underscoring the need to identify additional disease mechanisms and therapeutic targets.^18^ AML has been extensively studied at the genomic^19^, epigenomic^20^, transcriptomic, and proteomic levels^21, 22^, and recent large-scale proteomic studies have defined clinically relevant AML protein-expression states^23, 24, 25, 26^. However, because most proteomic analyses identify peptides by searching spectra against reference protein databases, they can systematically overlook cancer-associated proteins that are translated but not annotated in those references.^21, 22, 25^ Recent advances in transcriptome assembly and high-resolution mass spectrometry have created a powerful opportunity to integrate transcriptome sequencing with high-accuracy proteomics to systematically discover non-canonical proteins that were previously inaccessible.^7, 27, 28^ In addition, emerging functional evidence suggests that non-canonical proteoforms can contribute to leukemogenesis^29, 30^, but the extent to which AML cells recurrently express reference-unannotated cancer-specific proteoforms remains largely undefined.

Here, we asked whether primary AML samples express recurrent proteoforms that are absent from the current human reference proteome. We analyzed paired short-read RNA-sequencing and data-independent acquisition (DIA) mass spectrometry data from a large AML patient cohort^22^ using ProteomeGenerator2 (PG2)^14^, which integrates transcriptome assembly and ORF prediction with high-accuracy mass spectrometry to generate sample-specific protein databases and identify their associated proteoforms. We then developed a discovery funnel based on peptide mass spectral quality, absence from healthy CD34+ hematopoietic progenitor cell controls, recurrence across individual patient AML samples, low sequence similarity to annotated proteins, and independent support in deeply profiled AML cell line proteomes. This strategy identified recurrent AML-specific proteoforms, and nominated tumor-specific non-canonical disordered, alpha-helical, and membrane-associated proteoform classes for future functional and therapeutic studies.

## Results

### Study patient cohort

We analyzed paired RNA-sequencing and DIA mass spectrometry data^22^ from 123 primary AML samples that successfully passed the PG2 workflow (Fig. 1). The cohort included de novo, secondary, and therapy-related AML and spanned favorable, intermediate, and adverse ELN2017 risk groups. Detailed clinical and molecular characteristics are provided in Supplementary Data 1.

**Figure 1.**
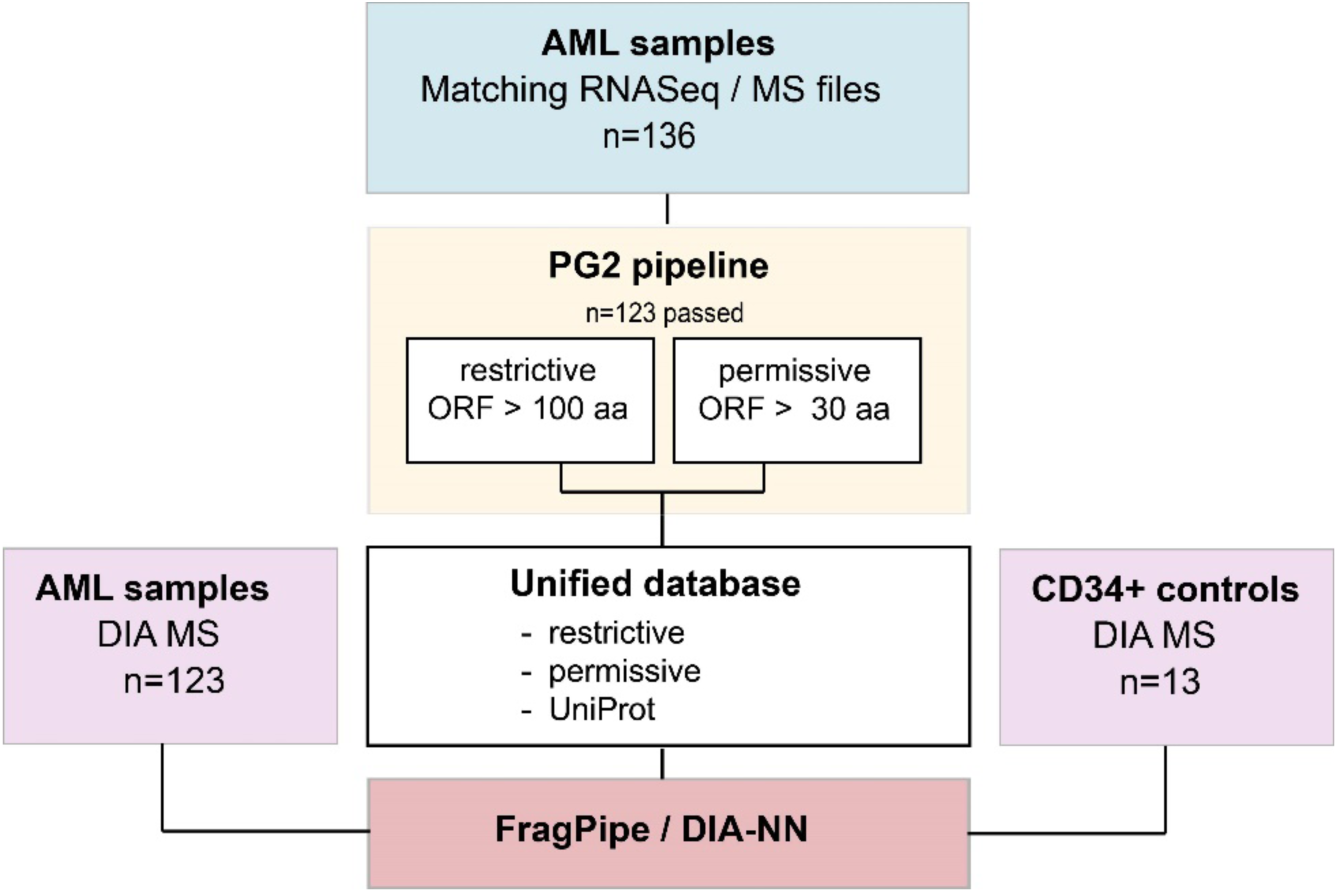
Proteogenomic workflow for discovery of AML-enriched reference-unannotated proteoforms. Paired RNA-sequencing and mass spectrometry data from 136 AML samples were processed with ProteomeGenerator2. A total of 123 samples passed the workflow. Restrictive and permissive ORF-prediction settings were used to generate sample-specific protein-sequence databases, which were combined with UniProt reference sequences to create a unified search database. DIA mass spectrometry data from AML samples and CD34+ hematopoietic controls were searched with FragPipe/DIA-NN. MS, mass spectrometry; DIA, data-independent acquisition. The figure was created with BioRender; https://BioRender.com/9fn3x77.

### Patient-specific proteogenomics reveals AML-specific reference-unannotated candidate proteoforms

Across all AML samples, we identified 13,265 Swiss-Prot-annotated protein isoforms, 9,682 protein groups, and 5,849 protein isoforms without Swiss-Prot annotation (Fig. 2a). Per patient sample, we identified a median of 5,420 protein isoforms, including a median of 5,093 (range 3669-6293) Swiss-Prot-annotated isoforms and 333 (137-644) reference-unannotated isoforms (Fig. 2b).

**Figure 2.**
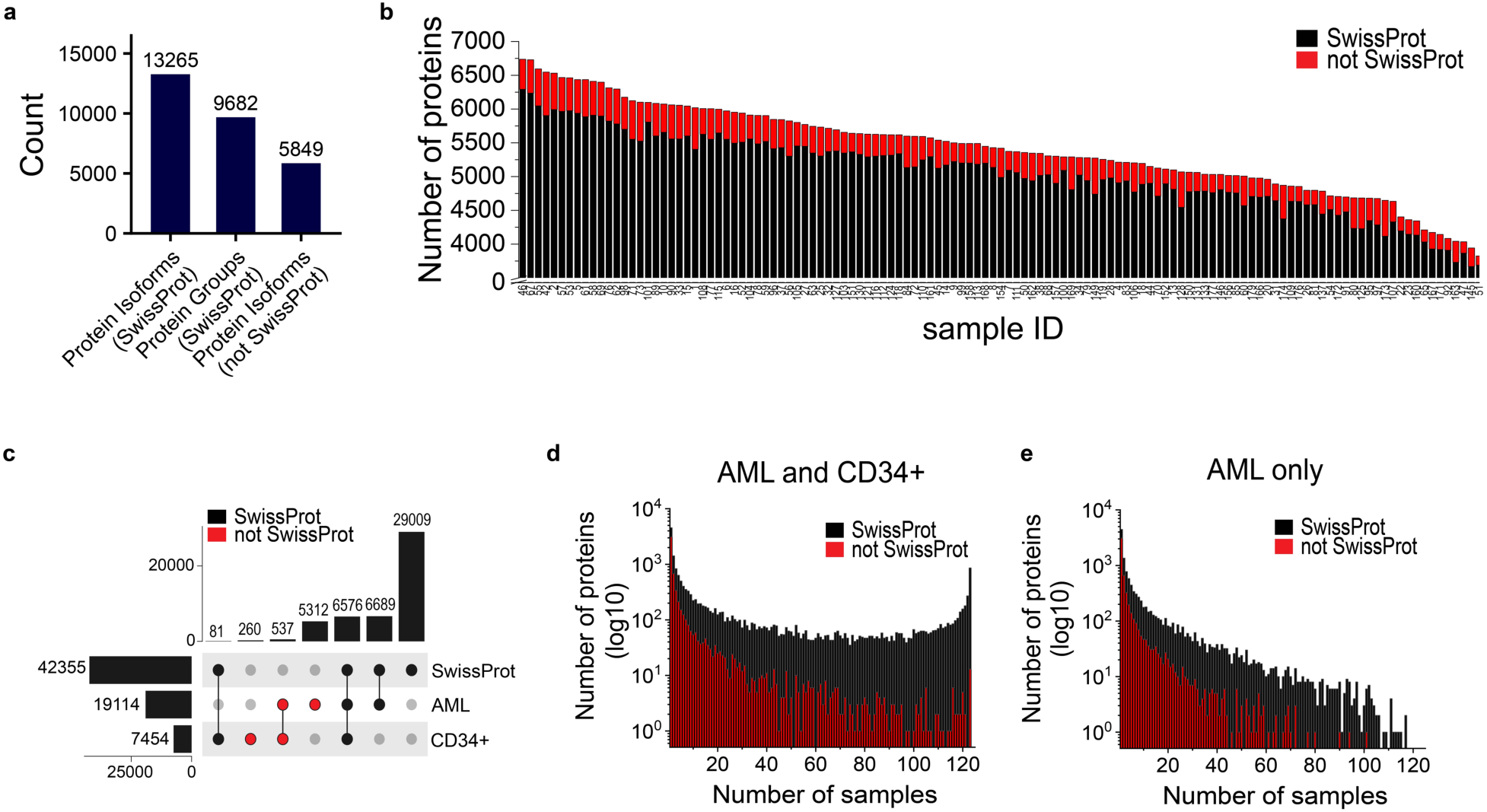
Patient-specific proteogenomics identifies reference-unannotated candidate proteoforms in AML. (a) Total numbers of identified Swiss-Prot–annotated protein isoforms, protein groups, and reference-unannotated candidate proteoforms across all AML samples. (b) Number of identified protein isoforms in individual AML samples, separated into Swiss-Prot-annotated isoforms and reference-unannotated candidates. (c) UpSet plot showing overlap among AML detections, CD34+ hematopoietic control detections, and Swiss-Prot annotation status. (d) Distribution of all detected protein isoforms according to the number of AML samples in which they were detected. The number of protein isoforms is shown on a log_10_ scale. (e) Distribution of AML-enriched protein isoforms after excluding detections present in CD34+ controls. The number of protein isoforms is shown on a log_10_ scale.

To identify AML-specific candidate proteoforms absent from normal hematopoietic progenitor cell controls, we searched mass spectrometry data from 13 CD34+ hematopoietic control samples^22^ using the same AML transcriptome-derived database. Of 13,265 Swiss-Prot annotated protein isoforms, almost half (6,576, 49.6%) were shared between AML and CD34+ samples. Most of the protein isoforms without Swiss-Prot annotation were detected in AML samples (5,312), and only few unannotated protein isoforms appeared shared between AML and healthy controls (537, Fig. 2c).

Most protein isoforms detected in only a single AML sample were reference-unannotated, whereas shared isoforms were predominantly annotated in Swiss-Prot. Specifically, among the 4,528 isoforms detected in individual AML samples, 3,127 lacked Swiss-Prot annotation and 1,401 were Swiss-Prot annotated (Fig. 2d). In contrast, among the 860 isoforms detected across all AML samples, 847 were Swiss-Prot annotated (Fig. 2d). After excluding protein isoforms detected in CD34+ hematopoietic controls, no AML-specific proteoforms were detected in every patient AML sample (Fig. 2e), consistent with varied biological subtypes and cells of origin known to occur among human AMLs.

### Analysis of leukemia-specific protein isoforms

We next focused on reference-unannotated candidate proteoforms detected specifically in AML samples and supported by at least one unique peptide. DIAMOND BLASTP^31^ was used to identify candidates with close similarity to annotated proteins. After retaining sequences with less than 80% identity to annotated proteins (Fig. 3a) and no significant match at e-value ≤1×10⁻³, 1,987 low-homology AML-specific candidate sequences remained. Of these, 39 were recurrently detected in more than 10% of AML patients (Fig. 3b), including 37 starting with N-terminal methionine (Supplementary Data 2).

**Figure 3.**
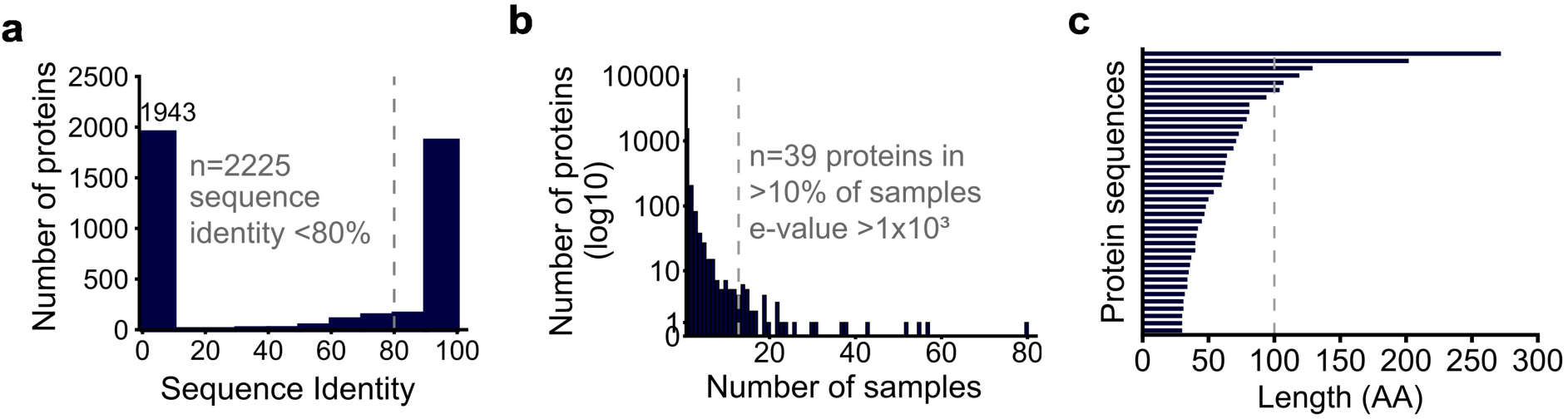
Recurrent AML-enriched candidate proteoforms lack close homology to annotated proteins. (a) Distribution of reference-unannotated candidate proteoforms in AML samples by percent amino-acid sequence identity to annotated proteins using DIAMOND BLASTP. The dashed line indicates the 80% sequence-identity threshold. (b) Number of low-homology candidate proteoforms detected in a given number of AML samples. The number of proteins is shown on a log_10_ scale. The dashed line indicates recurrence in more than 10% of AML samples; these candidates also lacked significant DIAMOND BLASTP matches to annotated proteins at e-value ≤1×10⁻³. (c) Amino-acid lengths of recurrent low-homology protein sequences detected only in AML samples. The dashed line indicates the 100-amino-acid threshold used to define microproteins.

Most recurrent candidates were relatively small in length: 33 of 39 predicted proteins were shorter than 100 amino acids (Fig. 3c). Each candidate was supported by at least one unique tryptic peptide of at least eight amino acids in length (Supplementary Fig. S2a). None of the supporting peptides had exact matches to annotated human protein isoforms in NCBI BLASTP^32^ or PeptideAtlas^33^. Mass-spectral assignments were highly confident, with PEP values below 0.01 for all but one supporting peptide, and quantified peptide intensities ranging between 1.3×10⁶ and 5.3×10⁸ across candidates (Supplementary Fig. S2b). Because most candidates were microproteins and therefore contained few protease-accessible peptides, we considered one high-confidence unique peptide sufficient for nomination but not for definitive functional annotation.^5^ One 272-amino-acid candidate was supported by two non-nested unique peptides of 14 and 24 amino acids (Supplementary Data 2). Consistent with the high accuracy of this analysis, detected peptides corresponding to this unannotated proteoform were also recently reported in an independent proteogenomic analysis of pediatric and adult AML^27^.

### Deep multi-protease DIA proteomics provides independent support for AML candidate proteoforms

To seek independent proteomic support for candidate proteoforms identified in primary patient AML samples, we performed deep DIA proteomics analysis of five AML cell lines and two CD34+ hematopoietic control samples using five different proteases, high-pH reversed-phase fractionation, and dia-PASEF acquisition using timsTOF Ultra mass spectrometer (Fig. 4a). We first searched the data against the UniProt reference database. This deep proteomic analysis identified 19,692 reference-annotated protein isoforms, including 14,181 isoforms supported by at least one unique peptide, corresponding to 12,266 protein groups (Fig. 4b). We detected 6,500 protein isoforms uniquely present in AML cell lines, and 11,913 (60.5%) protein isoforms shared between AML and CD34+ controls (Fig. 4c).

**Figure 4.**
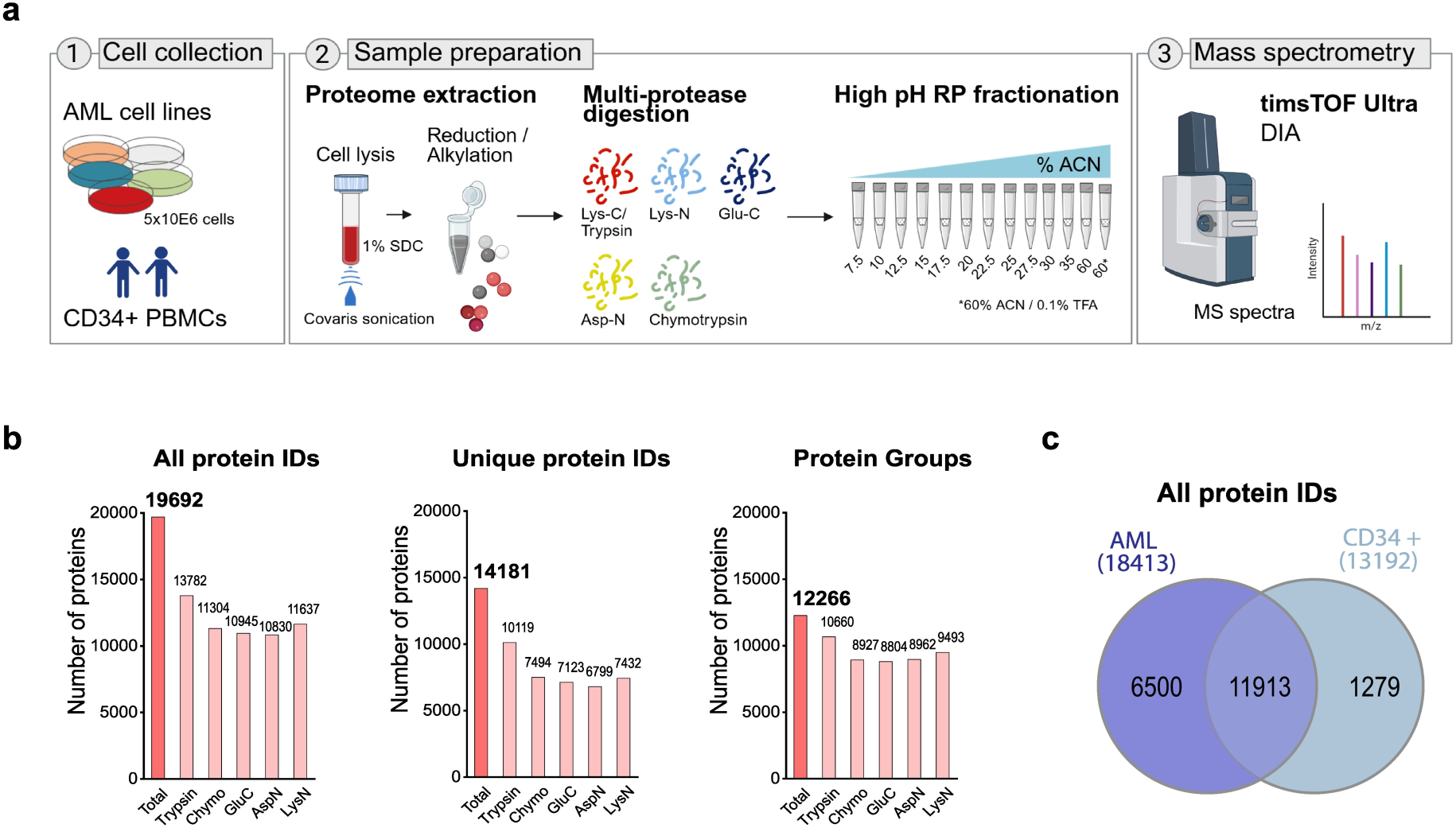
Deep multi-protease DIA proteomics provides orthogonal support for candidate proteoforms. (a) Workflow for deep proteomic profiling of five AML cell lines and two CD34+ hematopoietic controls. Whole-cell lysates were digested with five proteases, fractionated by high-pH reversed-phase chromatography, and analyzed by dia-PASEF mass spectrometry. The figure was created with BioRender; https://BioRender.com/cxgqew7. (b) Numbers of identified protein isoforms, unique protein isoforms, and protein groups for each protease and across the combined multi-protease dataset. (c) Overlap of protein isoforms detected in AML cell lines and CD34+ hematopoietic controls. SDC, sodium deoxycholate; ACN, acetonitrile; TFA, trifluoroacetic acid.

Comparative analysis of the cell line DIA data against the unified primary AML transcriptome-derived proteome database identified 404 reference-unannotated protein sequences detected in both primary AML samples and AML cell lines. Of these, 14 had less than 80% sequence identity to annotated proteins and no significant DIAMOND BLASTP^31^ matches at e-value ≤1×10⁻³ (Fig. 5a and 5b). Nearly all of these independently validated AML-specific proteoforms were microproteins: 13 of 14 were shorter than 100 amino acids, whereas only one proteoform exceeded this threshold (Fig. 5c). All protein sequences were supported by at least one unique peptide with a minimum length of 8 amino acids (Supplementary Fig. S3a). One 133-amino-acid candidate was supported by two non-nested unique peptides (Supplementary Data 2). Quantified unique peptides were detected for 13 candidates, and were relatively abundant, with average intensities ranging from 2.6×10⁴ to 1.2×10⁹. All supporting peptides showed high-confidence spectral assignments in both AML patient samples and AML cell lines, with PEP values below 0.01 (Supplementary Fig. S3b and S3c).

**Figure 5.**
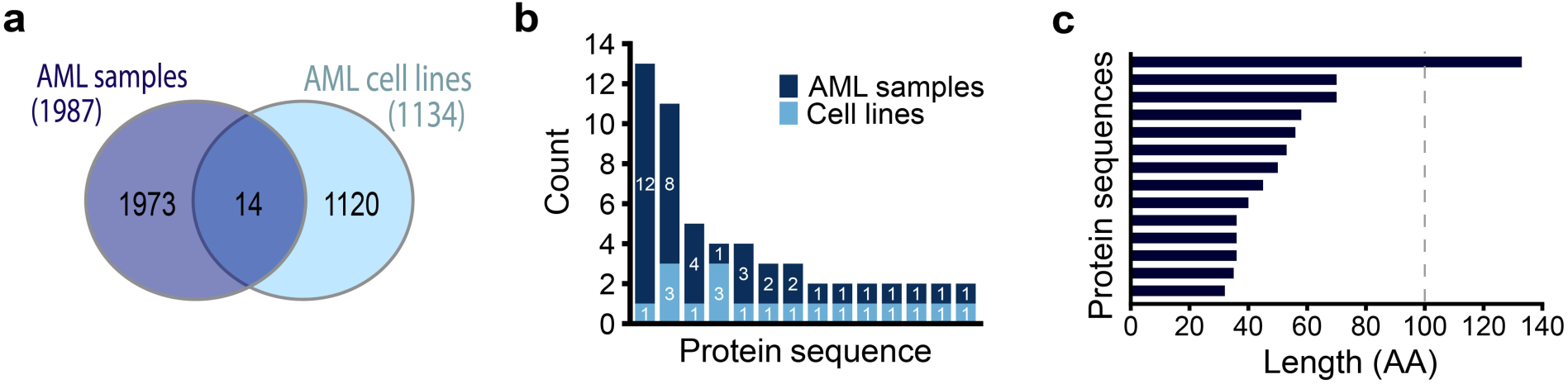
Candidate proteoforms detected in both AML primary samples and cell lines. (a) Overlap of low-homology reference-unannotated candidate proteoforms detected in primary AML samples and AML cell lines. Candidates were retained if they had less than 80% sequence identity to annotated proteins and no significant DIAMOND BLASTP match at e-value ≤1×10⁻³. (b) Recurrence of shared low-homology candidates across AML samples and AML cell lines. (c) Amino-acid lengths of shared candidate proteoforms. The dashed line indicates the 100-amino-acid threshold used to define microproteins.

### Predicted structural classes of AML-enriched candidate microproteins

To explore the structural and functional features of AML-specific proteoforms, we used InterPro^34^, including Pfam domain and family annotations, to analyze a combined set of 53 proteoforms comprising those recurrently detected in more than 10% of the patient samples and those detected overlapping in both AML cell lines and patient samples. Twenty-six proteoforms showed no predicted domain and structure annotations, 15 contained predicted intrinsically disordered regions, five contained predicted transmembrane regions, four contained annotated domains, and three contained signal peptides (Fig. 6a).

**Figure 6.**
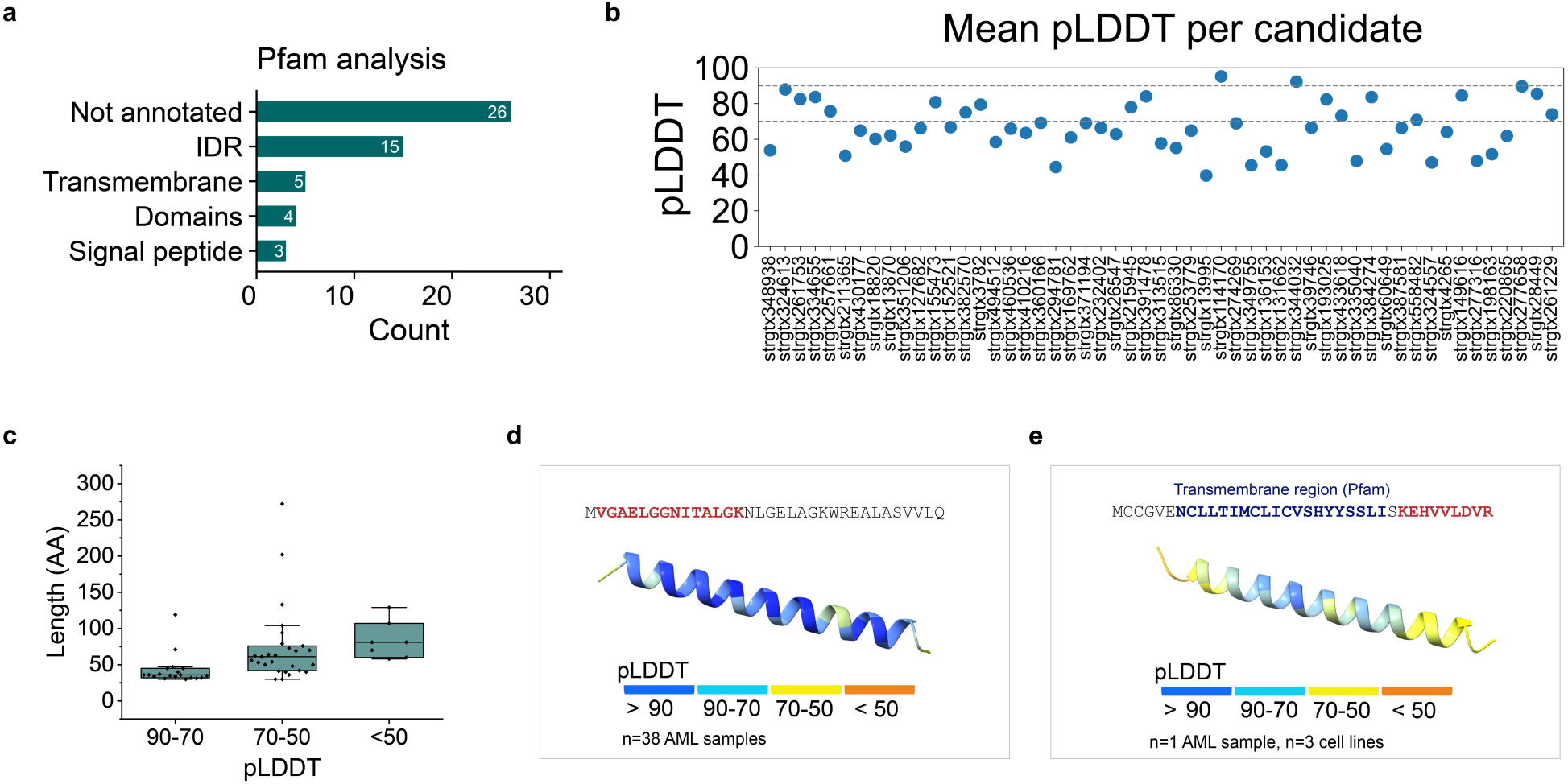
Predicted structural classes of recurrent AML-enriched candidate proteoforms. (a) InterPro/Pfam annotation of recurrent or cell line-supported low-homology candidate proteoforms. Bars indicate the number of candidates with each predicted feature. IDR, intrinsically disordered region. (b) Mean AlphaFold 3 pLDDT values for each candidate protein. (c) Relationship between candidate length and AlphaFold 3 model confidence, grouped by high, intermediate, and low mean pLDDT values. (d) AlphaFold 3 model of a recurrent alpha-helical microprotein detected in 38 AML samples. The mass spectrometry detected peptide is highlighted in red. (e) AlphaFold 3 model of a candidate microprotein detected in one AML sample and three AML cell lines. The supporting peptide is highlighted in red, and the predicted transmembrane region is highlighted in blue.

AlphaFold 3^35^ produced high-confidence structural models, defined by mean predicted local distance difference test (pLDDT) values greater than 70 for 19 candidates (Fig. 6b, Supplementary Fig. S4a and S4b). The remaining 34 candidates had lower-confidence structural models, including 11 protein sequences with predicted intrinsically disordered regions (Supplementary Fig. S5a and S5b). Higher-confidence structural models were enriched among shorter proteoforms (Fig. 6c) and were predominantly alpha-helical in structure (Fig. 6d). Three high-confidence alpha-helical proteoforms also contained predicted transmembrane regions (Fig. 6e). Together, these analyses defined a recurrent AML-specific non-canonical proteome composed of proteoforms that are absent from healthy CD34+ hematopoietic controls, divergent from annotated human proteins, and in a subset of cases, independently confirmed in deeply profiled human AML cell lines.

## Discussion

This study expands the AML proteome beyond the boundaries of current reference databases. Standard proteomic analyses identify peptides by searching spectra against annotated protein sequences; consequently, translated products absent from those databases can remain undetected. This limitation is particularly relevant in cancer, where somatic variation, aberrant transcription and splicing, and altered translation can generate proteoforms that are not represented in curated references. The Human Proteome Organization-Human Proteome Project^36^ and the TransCODE Consortium^5, 8^ have emphasized the need for rigorous criteria to annotate proteins encoded by non-coding or small ORFs. Our results apply this challenge to AML and show that patient leukemias recurrently express proteoforms missing from the current human reference proteome.

Using patient-specific proteogenomics, healthy CD34+ hematopoietic controls, mass spectral qualification, recurrence analysis, homology exclusion, and independent deep cell line proteomics, we identified a set of AML-specific reference-unannotated proteoforms. Thirty-nine unannotated proteoforms were detected in more than 10% patient leukemias studied, and 14 were also supported by independent multi-protease high-fractionation dia-PASEF mass spectrometry proteomics in AML cell lines. These findings indicate that AML contains a recurrent cryptic proteome that is systematically undetected when mass spectrometry data are searched only against standard references.

The leukemia-specific reference-unannotated proteoforms were enriched for small proteins, many of which were predicted to contain intrinsically disordered regions. This feature is biologically plausible because intrinsically disordered microproteins can act through short linear motifs, transient binding interfaces, and modulation of larger protein complexes.^2^ For example, NBDY, a 68-amino-acid microprotein that regulates the mRNA decapping complex, illustrates how a small disordered protein can control a larger ribonucleoprotein assembly.^37, 38^ In AML, the APPLE microprotein has been reported to promote leukemic cell proliferation and regulate oncogenic translation.^30^ Although the non-canonical proteoforms identified here require further direct functional studies, their predicted structural features suggest that some may act as regulators of protein interactions rather than enzymes or scaffolding proteins.

In addition to the identified intrinsically disordered microproteins, structural modeling using AlphaFold 3 identified a second class of microproteins with predominantly alpha-helical conformations^2^. Several of these proteoforms also contained predicted transmembrane domains, suggesting possible membrane-associated functions. Experimentally characterized alpha-helical microproteins can regulate diverse cellular processes, including regeneration^39^ and signaling^40, 41^. In cancer, alpha-helical transmembrane microproteins such as SMIM26^29, 42^ and SMIM30^43^ have been linked to malignant cell phenotypes, supporting the plausibility that some of the identified leukemia-specific proteoforms may contribute to AML biology, although their specific functions will require dedicated functional studies that are beyond the scope of the current analysis.

The structural features of identified AML-enriched reference-unannotated proteoforms suggest several testable models for how they could contribute to leukemia biology. Intrinsically disordered microproteins may function as short interaction modules that alter transcriptional, translational, or ribonucleoprotein complexes; alpha-helical microproteins may form compact binding elements or dominant modulators of larger protein assemblies; and candidates with transmembrane regions or signal peptides may localize to secretory, organellar, or cell surface compartments. Because human AML reflects aberrant hematopoietic cell differentiation and altered transcriptional and translational states, these proteoforms may report or regulate malignant cell states that are not captured by genetic analyses alone. Thus, the AML dark proteome may provide both mechanistic insights into leukemia biology and a source of candidate tumor-selective therapeutic targets.

Several limitations of the current study point directly to future work. Because most detected proteoforms were supported by single unique peptides, targeted proteomics with synthetic standards, orthogonal fragmentation methods, and endogenous tagging will be needed to confirm protein expression, abundance, and localization. Similarly, because absence from healthy CD34+ controls established AML enrichment but not absolute tumor specificity, larger studies across normal hematopoietic states and diverse healthy tissues will be required to define tumor selectivity. Lastly, this study relied on short-read sequencing for transcriptome assembly, and future studies can leverage long-read sequencing to improve both the specificity and sensitivity of transcriptome assembly, thereby improving resolution of splice isoforms, retained introns, fusion transcripts, and non-canonical and small ORFs, and expanding the discovery of cryptic AML proteoforms.

Although their functions remain to be established experimentally, results presented here identify recurrent AML-enriched proteoforms that are absent from standard reference databases and therefore nominate a prioritized set of hidden leukemia proteoforms. These findings also define priorities for improved dark-proteome discovery. Long-read RNA sequencing can resolve transcript structures; ribosome profiling can provide orthogonal evidence of translation; targeted proteomics can verify specific protein sequences and quantify proteoforms across larger cohorts; epitope tagging and CRISPR perturbation can test protein expression and function; and immunopeptidomics and cell surface proteomics can determine whether candidate microproteins generate leukemia-associated antigens that can be targeted therapeutically. Notably, dark proteome discovery is not restricted to unannotated ORFs. Recent evidence indicates that canonical transcripts can also generate stable, non-genetically-encoded proteoforms through alternate RNA decoding, which can be highly abundant compared to the canonical proteins and be selectively expressed in cancer tissues.^44^ Together, these approaches should reveal cryptic tumor-specific proteoforms that contribute to cancer pathogenesis and create new opportunities for biomarker discovery, immune recognition, and therapeutic targeting.

## Methods

### Generation of sample-specific databases for proteogenomics analysis

Raw RNA-sequencing, DNA-sequencing, and mass spectrometry data from the AML cohort reported by Jayavelu et al.^22^ were obtained from EGAD00001008484, EGAD00001008501, EGAD00001008485, and PXD023201. Mass spectrometry data for CD34+ hematopoietic controls were obtained from PXD028007^22^. PG2^14^ was run separately for each AML sample using matched DNA- and RNA-sequencing data aligned to GRCh38 with GENCODE v31 annotation. De novo transcriptome assembly was performed with StringTie^45^, and open-reading-frame prediction was performed with TransDecoder^46^. The minimum predicted ORF length was set to 100 amino acids in the restrictive run and 30 amino acids in the permissive run to enable detection of small protein candidates. To determine the transcriptomic origins of the protein sequences, we took the haplotype-specific gtf files from PG2 and used gffcompare (version 0.12.6) to match transcripts to GENCODE v45 based on intron structure. Using the gffcompare results, we reannotated each protein in the PG2 proteomes with a custom python script (https://github.com/kentsislab/fragpipe-dia/tree/pg2headers) with its corresponding transcript and gene from GENCODE v45.

Predicted protein sequences from the permissive and restrictive PG2 calculations from all AML samples were combined with UniProt UP000005640 (canonical and isoform sequences, downloaded on 02-06-2025) using tcdo-pg-tools version 0.1.0 (https://github.com/shahcompbio/tcdo_pg_tools). To control the search space created by private sample-specific predictions, we retained predicted non-reference sequences present in at least three AML transcriptomes before mass spectrometry searching. The resulting unified database contained 2,617,928 protein sequences (Supplementary Fig. S1) and was used for FragPipe/DIA-NN searches.

A total of 123 AML samples successfully passed the pipeline and were used for subsequent analysis. Clinical and molecular characteristics as provided by the original publications are given in the Supplementary Data 1.

### Generation of comprehensive proteomes from cell lines

#### Cell culture

The human AML cell lines Kasumi-1, MV4-11, U937, OCI-AML3 and OCI-AML2 were cultured at 37 °C with 5% CO_2_ in RPMI-1640 medium (Corning) supplemented with 10% fetal bovine serum (FBS, Gibco), 1% L-glutamine and 100 U/ml penicillin and 100 μg/ml streptomycin (Corning). Cell lines were authenticated by STR genotyping through the Integrated Genomics Operation, Center for Molecular Oncology at MSKCC and the absence of Mycoplasma species contamination was verified with the MycoAlert Mycoplasma detection kit (Lonza).

CD34+ cells from cryopreserved G-CSF-mobilized peripheral blood stem/progenitor cell products were obtained from the MSKCC biobank under IRB protocol #17-387 and from the Fred Hutch Cancer Center under protocol #1362.00. CD34+ cells were enriched using the Miltenyi CliniMACS system or the human CD34 MicroBead Kit (Miltenyi, #130-046-702) according to the manufacturer’s instructions.

#### Whole-cell proteomics

Five million cells from AML cell lines and CD34+ enriched cells were washed with PBS once and pelleted at 300 × *g* for 5 min at 4 °C. The supernatant was completely removed and the cell pellet was snap frozen on dry ice before storage at −80 °C.

For proteome extraction, cells were resuspended in lysis buffer containing 1% sodium deoxycholate (Sigma-Aldrich), 50 mM ammonium bicarbonate ∼pH 8.0 (Sigma-Aldrich), 1x PhosSTOP (Roche) and 1x cOmplete™ EDTA-free Protease Inhibitor Cocktail (Roche). The cell pellet was lysed by sonication (Covaris S220) at 4 °C for 6 min (peak power 175 W, Duty factor 10%, Cycles/burst 200). Lysate protein concentration was quantified using the Detergent Compatible Bradford Assay Kit (Pierce). Protein was reduced using 10 mM dithiothreitol (DTT) for 30 min at room temperature and alkylated using 25 mM iodoacetamide (IAA) in the dark for 45 min. The protein lysate volume was equally distributed to new maximum recovery tubes (Axygen) for each digestion reaction. Digestion with LysC/trypsin (Promega, V507A), GluC (Promega, V165A) and rAspN (Promega, VA116A) was performed using a total 1:50 protease/proteome ratio (2/3 protease at 37 °C overnight, 1/3 protease for 2 h at 37 °C). Chymotrypsin (Promega, V106A) digestion was carried out using a 1:80 protease/proteome ratio at 25 °C (2/3 protease at 25 °C overnight, 1/3 protease for 2 h at 25 °C). Digestion with LysN (AVS Bio Netherlands B.V.) was performed using a 1:20 protease-to-proteome ratio (2/3 protease at 37 °C overnight, 1/3 protease for 2 h at 37 °C). Reactions were stored at −80 °C until further use.

High-pH reversed-phase (HpH-RP) fractionation was manually performed using the high-pH RP fractionation kit (Pierce, #84868). Samples were acidified with 5% formic acid (FA), incubated for 10 min at room temperature, and debris removed with centrifugation at 18000 × *g* for 10 min at 4 °C. BioPureSPN MACRO solid phase extraction (SPE) columns (Nest Group, HMM S18R) were washed with 100% acetonitrile (ACN, Fisher Chemical) and equilibrated 2 times with 0.1% Trifluoroacetic acid (TFA, Thermo Scientific). The cleared sample supernatant was loaded on the column and centrifuged at 60 × *g* until completely passed through the column and the flow-through collected in a separate tube. The column was washed 3 times with MS-grade water (Fisher Chemical) and the flow-through pooled together. Eluates of the individual fractions (in total n=13) were collected in separate low-binding tubes (Axygen) using increasing ACN concentrations in 0.1% triethylamine. Fractions 1 to 10 ranged from 7.5% ACN to 30% ACN with increasing steps of 2.5% ACN per fraction (fraction 1 was eluted 2 times), followed by 35% and 60% ACN elution and a final elution step using 60% ACN/0.1% TFA (2 times). The flow-through was desalted using BioPureSPN MIDI SPE columns (Nest Group, HEM S18R). Columns were washed with 100% ACN and equilibrated 2 times with 0.1% TFA. After sample loading, the column was washed 2 times with 0.1% TFA before elution using stepwise 50% ACN/0.1% TFA and 60% ACN/0.1% TFA. Fractions were lyophilized and stored at −80 °C until further use.

#### LC-MS

Liquid chromatography-mass spectrometry (LC-MS) analysis of the AML cell lines and CD34+ controls was performed using the nanoElute 2 system (Bruker Switzerland AG) coupled to the timsTOF Ultra 2 mass spectrometer (Bruker Daltonics GmbH & Co.). Peptide samples (∼100 ng in 1 µL) were directly loaded onto a 25 cm C18 reverse-phase column (75 µm inner diameter, 1.5 µm particle size; PepSep Ultra, Bruker Daltonics GmbH & Co.). Chromatographic separation was carried out at a flow rate of 300 nL/min using a linear gradient from 3% to 22% buffer B (99.9% acetonitrile, 0.1% formic acid) over 25 minutes, then an increase to 30% buffer B over 5 min, followed by an increase to 95% buffer B sustained for 8 minutes (total run time: 38 minutes). The column temperature was maintained at 50 °C using an integrated column heater. Data-independent acquisition using parallel accumulation-serial fragmentation (dia-PASEF) was employed, utilizing 25 isolation windows. Each window had a width of 26 Da with a 1 Da overlap and was configured with variable ion mobility (IM) settings. Two acquisition steps were used (1 MS1 ramp and 11 MS/MS ramps), yielding a total cycle time of 0.65 seconds. The m/z range was set from 299.5 to 1200.5, and the ion mobility range (1/K₀) spanned from 0.70 to 1.30 V·s/cm². Ramp and accumulation times were both set to 75.0 ms. Collision energy was dynamically adjusted from 20 to 63 eV across the ion mobility range (1/K₀) from 0.60 to 1.60 V·s/cm².

#### FragPipe analysis

DIA AML patient data were analyzed using FragPipe (version 23.0) with the DIA_SpecLib_Quant workflow and peptide quantification performed with DIA-NN (version 2.1.0) at 1% FDR. diaPASEF data acquired on the timsTOF instrument from the cell lines were analyzed using FragPipe version 22.0 using the DIA_SpecLib_Quant_diaPASEF workflow and peptide quantification was performed with DIA-NN (version 1.8.2) at 1% FDR.

In diaTracer, “Delta Apex IM” was set to 0.01, “Delta Apex RT” was set to 3, “RF max” was set to 500, and “Corr threshold” was set to 0.3. The mass defect filter was enabled. In both workflows, the respective protease was selected in MS Fragger (“stricttrypsin”, “aspn”, “chymotrypsin”, “gluc”, “lysn”) and 2 missed cleavages allowed. The peptide length was set to 7-50. The initial precursor and fragment mass tolerances were set to 10 ppm and 20 ppm, respectively. Spectrum deisotoping, mass calibration, and parameter optimization were enabled. Oxidation of methionine and N-terminal acetylation were set as variable modifications. Carbamidomethylation of cysteine was set as a fixed modification. The unified transcriptome database was appended with common contaminants and decoys and used in the searches. In addition, the multi-protease MS data generated from cell lines and CD34+ controls were searched against the UniProt UP000005640 canonical and isoform database (downloaded 01-24-2025, appended with common contaminants and decoys). AML patient samples were analyzed individually. AML cell lines were analyzed individually (per cell line and enzyme) with all fractions set as consecutive experiments. All fractions from n=2 CD34+ control samples were analyzed together for each enzyme individually.

#### Data analysis

Predicted protein sequences were collapsed across samples by exact amino-acid sequence identity. Identical sequences detected in multiple samples were assigned a shared candidate identifier, and sample recurrence was calculated from the number of AML samples in which the corresponding sequence was present in the PG2-derived database and supported by mass spectrometry peptide evidence.

Sequence similarity to annotated proteins was assessed with DIAMOND BLASTP^31^ (version 2.1.12) against the UniProt UP000005640 database (canonical and isoform sequences, downloaded 03-25-2026). Candidates were considered to lack close homology to annotated proteins if they had less than 80% amino-acid sequence identity (pident) and no significant match at e-value ≤1×10⁻³. Peptide uniqueness was assessed by searching peptide sequences against NCBI BLASTP^32^ (accessed 08-26-2026) using the clustered non-redundant protein database (last update 07-12-2026), and against PeptideAtlas^33^ (Human 2026-01) using the online version of ProteoMapper^47^, which is an integrated component of the Trans-Proteomic Pipeline (version 5.2.0). Peptides with exact matches to annotated human proteins were excluded from the high-confidence candidate set.

Protein family similarities and structures were examined using Pfam in InterPro (version 109.0).^34^ Protein folding was modeled with AlphaFold 3 using the AlphaFold Server^35^ and visualized in UCSF ChimeraX (version 1.12)^48^. The pLDDT profiles and mean pLDDT were calculated using the “atom_plddt” per residue output and visualization generated with Python (version 3.12.4). Venn diagrams were generated using the InteractiVenn online tool.^49^ Figures were generated with OriginPro (2022b), R Studio (R version 4.6.0) and BioRender. Figures were edited with Adobe Illustrator 2026.

#### Peptide intensities and PEP score analysis

From the DIA-NN report output files, peptide precursor intensities were summed across charge states and fractions (AML cell lines) within each sample and the minimum posterior error probability (PEP) value was extracted. For recurrent peptides across samples, the mean peptide intensity (log10 transformed) across samples was used for visualization and the minimum observed PEP value (-log10 transformed) was retained as a confidence metric.

#### Data and code availability

All processed candidate tables, peptide-level evidence, and source data used for figure generation are available through Zenodo at 10.5281/zenodo.22127938. Raw mass spectrometry data generated in this study have been deposited to MassIVE under project number MSV000102980. The PG2 analysis pipeline is available at https://github.com/cBio-MSKCC/ProteomeGenerator2/tree/B-101-985. The Snakemake pipelines for reannotation of PG2 proteomes and FragPipe analysis are available at https://github.com/kentsislab/fragpipe-dia. The code for combining protein fasta files across multiple samples is available at https://github.com/shahcompbio/tcdo_pg_tools. The Nextflow pipeline for running DIAMOND BLASTP searches is available at https://github.com/kentsislab/ORFology. Analysis code and figure-generation scripts are available via Zenodo at 10.5281/zenodo.22127938.

## Acknowledgements

We thank Helen Mueller and for critical discussions. We thank Nicholas D. Socci, Caitlin Jones, Krista Kazmierkiewicz and the MSK Bioinformatics Core Facility for assistance with PG2 implementation, transcriptome-derived FASTA generation, and computational workflow support.

## Author contributions

A.K. and L.K.S. conceptualized the study. A.K., L.K.S., and K.K. designed and optimized experimental methods. A.K., L.K.S., A.P.S., and A.M. designed and optimized bioinformatic methods. K.K., J.Z., G.C., and L.K.S. performed experiments. L.K.S., K.K., A.P.S., and G.C. analyzed data. A.K. and L.K.S. wrote the original manuscript, which was subsequently edited by all authors. The authors used ChatGPT to refine the text for clarity, which was subsequently directly reviewed and edited to ensure accuracy.

