## Supplementary Figures for "Recurrent non-canonical proteoforms in acute myeloid leukemia identified by integrative proteogenomics"

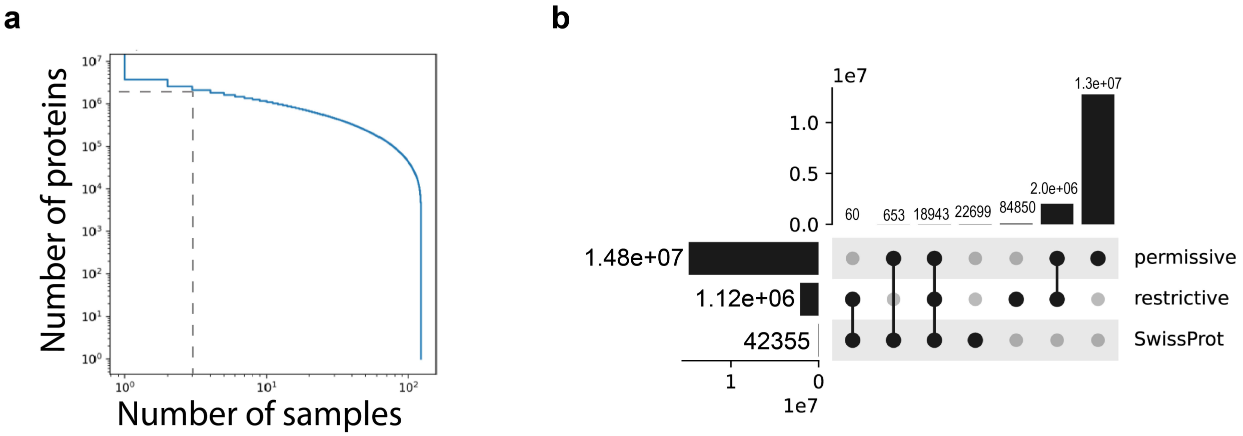


**Supplementary Figure S1. Construction of the unified transcriptome-derived protein database.** (a) Number of predicted protein sequences detected across AML transcriptomes before database inclusion. The dashed line indicates the minimum recurrence threshold of three AML samples used to retain predicted non-reference sequences for mass spectrometry searching. (b) UpSet plot showing overlap among UniProt reference sequences, restrictive PG2 predictions generated with ORF length greater than 100 amino acids, and permissive PG2 predictions generated with ORF length greater than 30 amino acids. The final unified database combined UniProt sequences with retained restrictive and permissive PG2 predictions and contained 2,617,928 protein sequences.

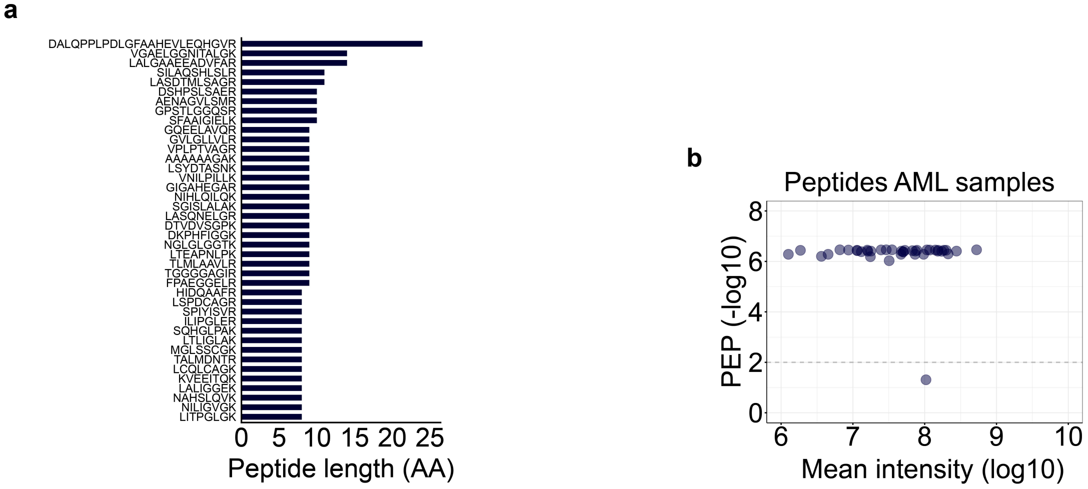


**Supplementary Figure S2. High-quality peptides supporting protein sequences recurrently detected in AML samples.** (a) Amino acid (AA) lengths of peptides supporting the 39 unannotated proteins recurrently detected in more than 10% of AML samples. (b) Minimum PEP scores (y-axis) and mean peptide intensities (x-axis) for quantified peptides supporting the 39 unannotated proteins recurrently detected in more than 10% of AML samples.

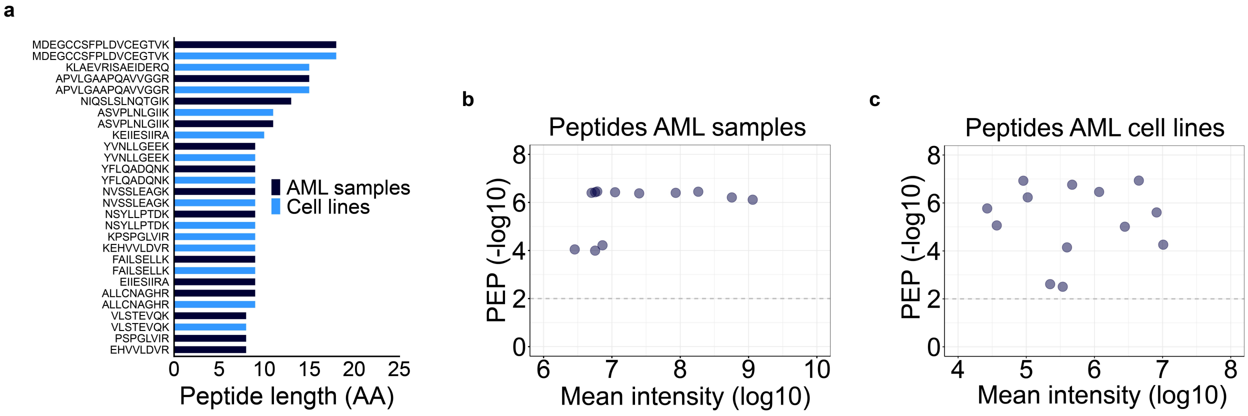


**Supplementary Figure S3. High-quality peptides supporting protein sequences detected in AML patient samples and AML cell lines**. (a) Amino acid (AA) lengths of peptides supporting the 14 unannotated proteins independently detected in both AML samples and AML cell lines. (b) Minimum PEP scores (y-axis) and mean peptide intensities (x-axis) for quantified peptides supporting the 14 unannotated proteins independently detected in both AML samples and AML cell lines. Values were derived from the AML patient dataset. (c) Minimum PEP scores (y-axis) and mean peptide intensities (x-axis; summed across fractions) for quantified peptides supporting the 14 unannotated proteins independently detected in both AML samples and AML cell lines. Values were derived from the AML cell line dataset.

**a**

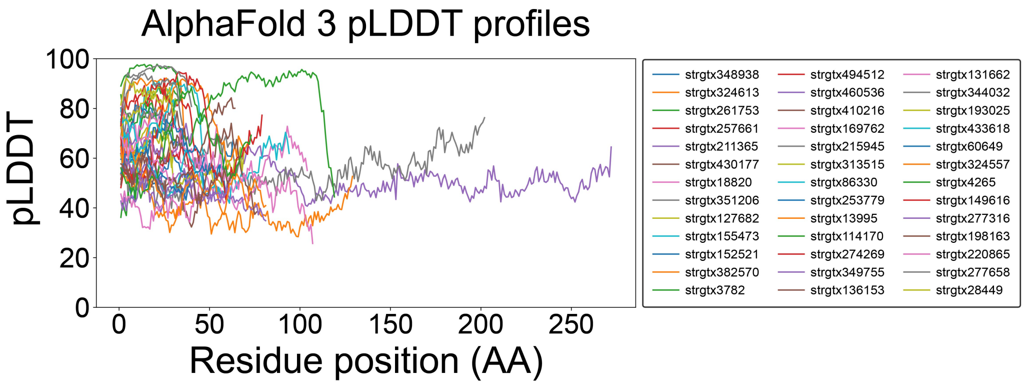


**b**

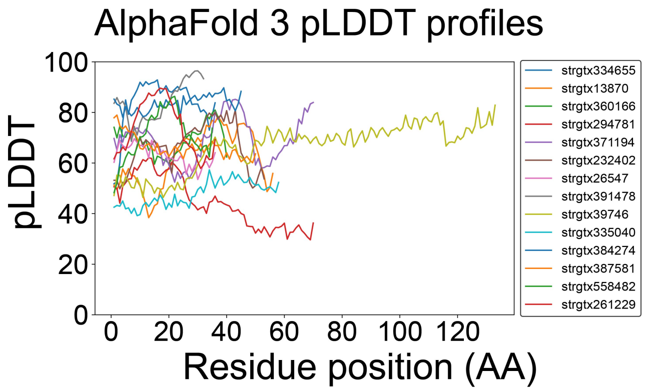


**Supplementary Figure S4. AlphaFold 3 pLDDT profiles per residue position of unannotated proteins detected in AML samples.** (a) pLDDT profiles from 39 unannotated proteins recurrently detected in >10% of the AML samples. The x-axis shows the residue position of the respective amino acid (AA) and the y-axis the respective confidence value given as pLDDT score. (b) pLDDT profiles from 14 unannotated proteins detected in AML samples and AML cell lines. The x-axis shows the residue position of the respective amino acid (AA) and the y-axis the respective confidence value given as pLDDT score.

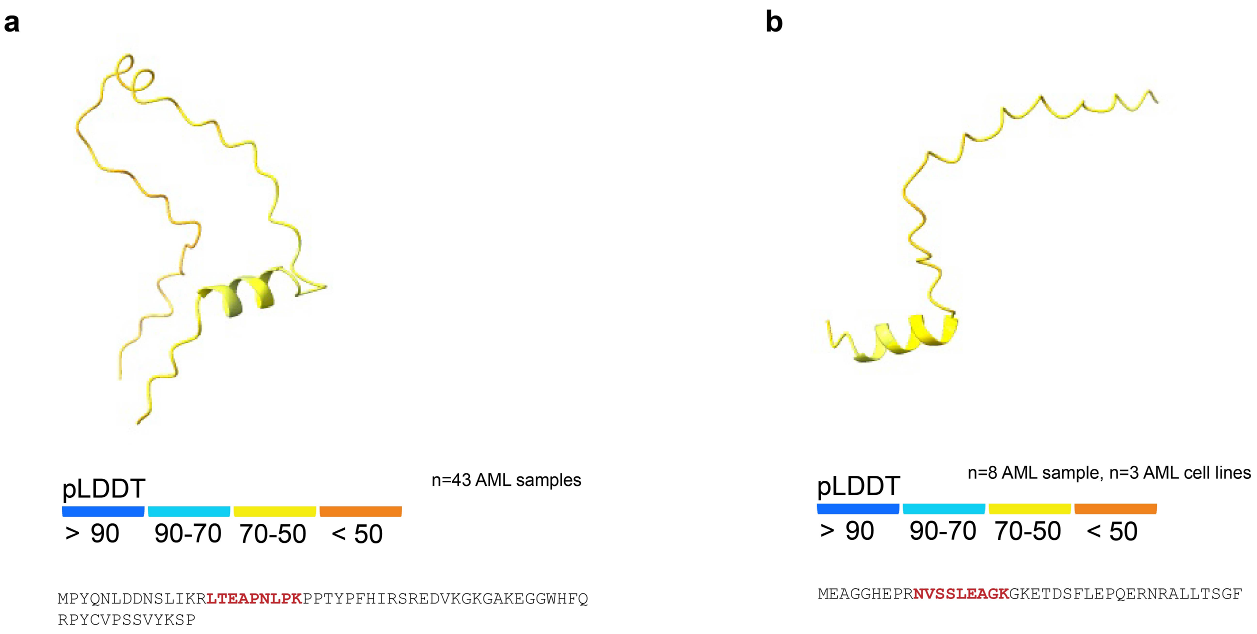


**Supplementary Figure S5. Examples of unannotated proteins with intrinsically disordered regions.** (a) Model protein folding (AlphaFold 3) of a small protein sequence with predicted IDR (Pfam) recurrently detected in 43 AML samples. Red highlights the supporting unique peptide in the protein sequence. (b) Model protein folding (AlphaFold 3) of a larger protein sequence with predicted IDR (Pfam) recurrently detected in 36 AML samples. Red highlights the supporting unique mass spectrometry detected peptides in the protein sequence.
